# Smooth muscle K_Ca_3.1 limits plaque progression and alters plaque composition in mice

**DOI:** 10.64898/2026.11.12.766210

**Authors:** P. Alam, D.L. Tharp, H.J. Bowles, H. Bui, S.B. Bender, D.K. Bowles

## Abstract

Increased expression of K_Ca_3.1 has been found in vascular smooth muscle cells (SMC), macrophages, and T cells in atherosclerotic lesions from humans and mice. While the efficacy of pharmacological inhibition and global silencing of K_Ca_3.1 in limiting atherosclerosis has been demonstrated, the relative cell-specific contributions of K_Ca_3.1 have not been determined. Therefore, we generated a smooth muscle (SMC) specific inducible K_Ca_3.1^-/-^ mouse with global Apoe^-/-^ knockout (iSMKO) and assessed lesion development in the brachiocephalic artery (BCA) of tamoxifen vs. vehicle injected iSMKO mice on a Western diet (WD) for 15 or 40 weeks. In BCAs of iSMKO mice, lesion size and relative stenosis were increased and lumenal area decreased compared to vehicle injected mice. Additionally, iSMKO mice exhibited a significant reduction in SMC and collagen content at 15 weeks and 40 weeks, while macrophage content increased at 40 weeks. RNAseq analysis of BCA samples from tamoxifen and vehicle injected iSMKO mice prior to WD feeding (baseline) indicated increased mitochondrial function and PPAR-dependent signaling, similar to that found in prior studies of global K_Ca_3.1 silencing. However, following 15 weeks of WD and induction of disease, these differences were no longer observed. Overall, this study provides the first cell-specific silencing of K_Ca_3.1 to elucidate the contribution of K_Ca_3.1 in smooth muscle to prior observations of K_Ca_3.1 silencing. Our findings reveal both a beneficial role of SMC K_Ca_3.1 in limiting plaque size and macrophage infiltration, and a potentially detrimental effect of reducing plaque SMC and collagen content. Combined with previous studies, this raises caution in the use of pharmacological K_Ca_3.1 inhibitors in atherosclerosis and highlights the need for cell-specific targeting of K_Ca_3.1 in optimizing potential therapies.

**Highlights:**

1. Smooth muscle specific genetic knockout of K_Ca_3.1 in mice results in larger atherosclerotic plaques with increased macrophage content revealing a beneficial role of smooth muscle K_Ca_3.1 in limiting atherosclerotic lesion development.
2. Silencing K_Ca_3.1 in smooth muscle leads to a significant reduction in smooth muscle cell and collagen content within the plaques, consistent with a role for K_Ca_3.1 in smooth muscle proliferation and medial-to-intimal cell migration during atherosclerosis.
3. Thus, our findings reveal both a beneficial role of SMC K_Ca_3.1 in plaque size and macrophage infiltration, and a potentially detrimental effect of reducing plaque SMC and collagen content, resulting in a lesion with characteristics of a more unstable plaque.
4. RNA sequencing predicted similar downstream signaling effects in both SMC-specific and global K_Ca_3.1 silencing, including increased mitochondrial function and PPAR signaling, which likely plays a role in inhibiting smooth muscle proliferation and medial-to-intimal migration in atherosclerosis.
5. While global inhibition of K_Ca_3.1 has been demonstrated to have beneficial effects on atherosclerosis development, this study highlights the need for cell-specific targeting of K_Ca_3.1 to optimize potential therapies.

## Introduction

Despite lipid-lowering and emerging anti-inflammatory agents, coronary heart disease remains a leading cause of death in both men and women in the United States ^1,2^. Over 20 million Americans > 20 years of age have coronary heart disease (CHD), and each year ∼635,000 Americans have a new coronary attack, and ∼300,000 have a recurrent attack^1^. Atherosclerosis is a chronic, inflammatory, and proliferative disease that develops over decades, involving multiple cell types, including endothelial cells, smooth muscle cells (SMCs), fibroblasts, macrophages, T-cells, B-cells, and platelets. The intermediate-conductance Ca^2+^ - activated K^+^ channel (K_Ca_3.1) is expressed in all of these cell types and plays a crucial role in T-cell, B-cell, fibroblast, and SMC proliferation, as well as the migration of SMCs, macrophages, and platelet coagulation^3–7^, leading to the consideration of K_Ca_3.1 modulators as potential therapies for vascular disease^8,9^.

Increased expression of K_Ca_3.1 has been found in atherosclerotic lesions from humans and mice^6,10^ and several studies have shown systemic delivery of K_Ca_3.1 inhibitors can attenuate atherosclerosis lesion development in mice^6,10–12^. We have recently provided genetic confirmation that global silencing of K_Ca_3.1 inhibits atherosclerotic lesion development ∼70% in both male and female mice ^13^. Together, these studies support the hypothesis that the upregulation of K_Ca_3.1 is a major contributor to SMC migration and proliferation during atherosclerosis and restenosis. However, the cell type-specific relative contribution of K_Ca_3.1 activation during atherosclerotic lesion development has not been determined. Substantial evidence in the literature indicates a role for SMC K_Ca_3.1 in lesion development and composition. Synthetic, proliferating SMC increase expression of K_Ca_3.1 to become the dominant K^+^ channel ^7,14^ and its upregulation has been observed in neointimal SMCs in balloon-injured rat carotid arteries^6,15^ as well as atherosclerotic lesions in mice and humans^6^. In addition, we have shown that acute administration of the K_Ca_3.1 inhibitor, TRAM-34, during coronary angioplasty in a swine model of coronary restenosis can inhibit lesion development, predominantly by targeting SMC proliferation ^4^. Migration of SMC from the media to the intima is a major contributor to both restenosis and atherosclerosis and we^4,7^ and others ^5,6^ have shown that K_Ca_3.1 is critical for SMC migration. Reduced lesion size due to global silencing of K_Ca_3.1 was also associated with a reduction in smooth muscle cell lesion content ^13^.

Thus, while there is substantial evidence supporting the role of K_Ca_3.1 in SMC contribution to atherosclerosis, evidence to date has been derived from *in vitro* studies or global pharmacological and genetic silencing. Direct evidence of the SMC-specific contribution to lesion development and composition is lacking. Therefore, the purpose of the current study was to provide the first conditional, SMC specific genetic silencing of K_Ca_3.1 in the context of atherosclerosis development *in vivo*.

## Material and Methods

### Ethics Statement

Experimental protocols performed adhered to ARRIVE guidelines, were approved by the University of Missouri Animal Care and Use Committee (Protocol #70602) and conducted in accordance with the “Principles for the Utilization and Care of Vertebrate Animals used in testing, Research and Training.” Animals were anesthetized with Ketamine/Xylazine and euthanized by exsanguination in accordance with American Veterinary Medical Association guidelines. Detailed methods can be found in Supplement.

### Statistical analysis

All data are presented as mean ± SE. Statistical analyses were performed with the Prism 10 statistical software (GraphPad Software, San Diego, CA). Before comparisons, normality was assessed using the Shapiro-Wilk method, and variance homogeneity was determined using F testing. Normal distributions with homogenous variances were compared using unpaired, 2-tailed Student t tests, one-way or two-way ANOVA as appropriate. A *P* value threshold of 0.05 was pre-established to designate statistical significance. Statistical cutoffs for differentially expressed genes were −1.0 ≤ log^2^FC ≥ 1.0 change in expression with an adjusted p < 0.05.

## Results

### Generation of SM-specific K_Ca_3.1, ApoE knockout mice

Inducible, smooth muscle specific knockout mice (iSMKO) were generated by CRISPR-mediate knockout of Apoe in Kcnn4^fl/fl^ mice and crossing of the resultant Kcnn4^fl/fl^Apoe^-/-^ mice with the Myh11-CreER^T^^2^ mice to generate Myh11-CreER^T^^2^ Kcnn4^fl/fl^Apoe^-/-^. Kcnn4^fl/fl^ mice were obtained from Kcnn4^fl/fl^ recombinant ES cell created by Biocytogen (Beijing, China) with LoxP sites flanking exon 4 of Kcnn4. F1 mouse generation and CRISPR-mediated Apoe knockout was performed by the University of Missouri Animal Modeling Core (Columbia, MO). To validate smooth muscle specificity of our Myh11-CreER^T^^2^ Kcnn4^fl/fl^Apoe^-/-^ (iSMKO), we crossed our mice with a mTmG reporter mice (#007676, Jackson Laboratory). Cre-mediated excision results in the replacement of red Tomato reporter with green GFP (Fig. 1a). Aortas from vehicle (Fig. 1b) or tamoxifen-injected (Fig 1c) mice on normal chow diet demonstrate Cre-excision exclusively in the media. Similarly, aortas from Western diet fed mice show medial and intimal Cre-excision in tamoxifen injected mice (Fig. 1e & g), but only minimal, non-specific autofluorescence in vehicle injected mice (Fig. 1d & f). Smooth muscle specificity is also shown in sections of myocardium of tamoxifen-injected mice (Fig. 1h) demonstrating GFP expression restricted to the media of a coronary artery. Furthermore, tamoxifen injection significantly reduced K_Ca_3.1 mRNA in the aorta (Fig. 1i). Importantly, no significant difference was observed in cholesterol levels between vehicle and tamoxifen-injected mice on a Western diet (1098 ± 36 vs 1163 ± 35 mg/dl, resp.).

**Figure 1.**
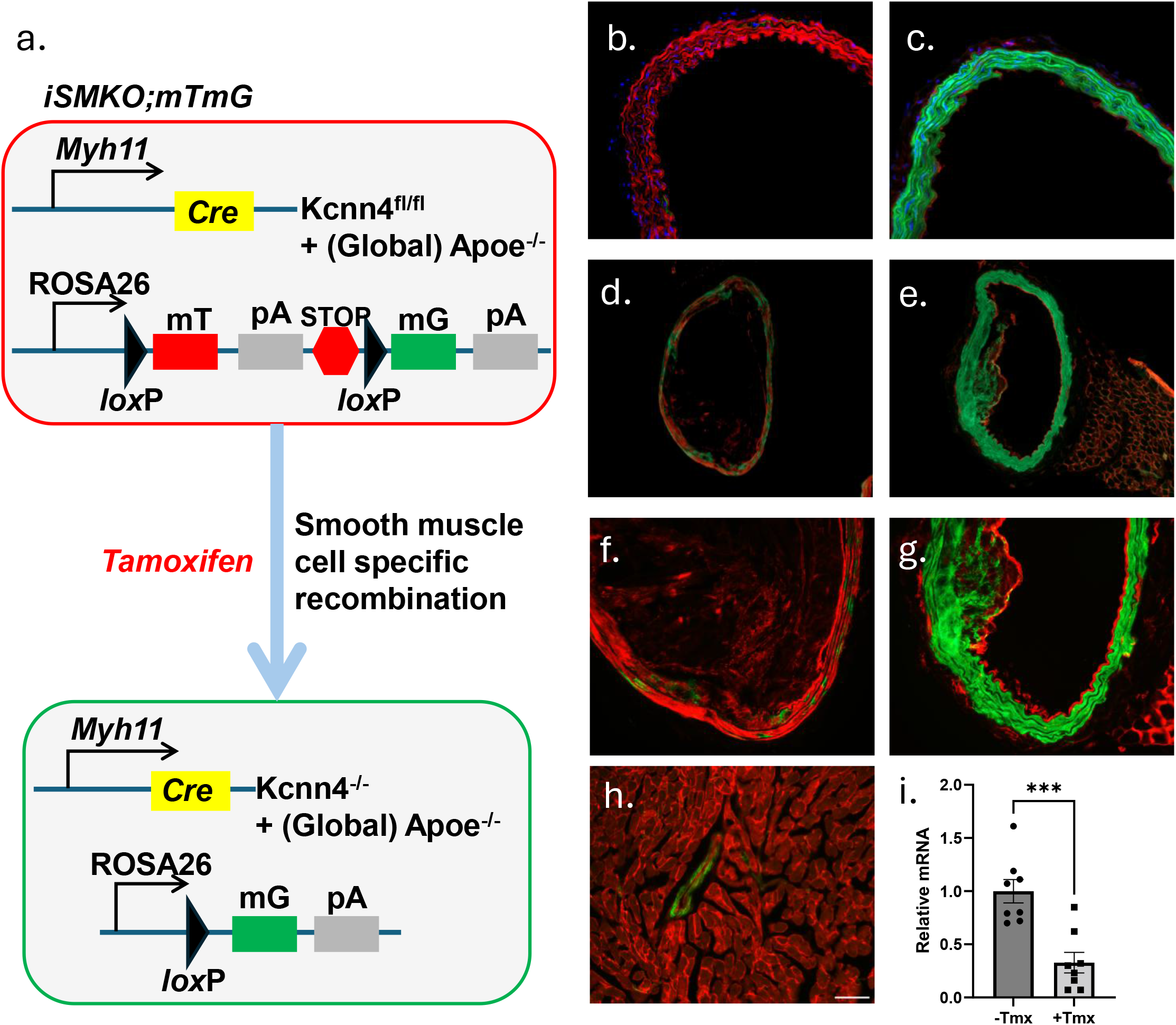
*Validation of Myh11-CreER^T2^ Kcnn4^fl/fl^Apoe^-/-^ (iSMKO) mice.* a.) Tamoxifen injection of iSMKO mice crossed with mTmG reporter mice results in Cre-mediated excision and replacement of red Tomato reporter with green GFP. Aortas from vehicle (b) or tamoxifen-injected (c) mice on normal chow diet demonstrate Cre-excision exclusively in the media. Aortas from Western diet fed mice show medial and intimal Cre-excision in tamoxifen injected mice (e & g), but only minimal, non-specific autofluorescence in vehicle injected mice (d & f). Myocardium of tamoxifen-injected mice (h) demonstrating GFP expression restricted to the media of a coronary artery. i.) Tamoxifen injection reduces K_Ca_3.1 mRNA in the aorta.-Tmx, vehicle (n=8); +Tmx, tamoxifen-injected, (n=8); ***p < 0.001.

### Brachiocephalic artery morphometry

We examined atherosclerosis development in the brachiocephalic artery (BCA) at both 15 and 40 weeks of WD (Fig. 2a-d). As expected, iSMKO mice fed a Western diet developed complex atherosclerotic lesions consisting of a fibrous, cellular matrix overlying lipid cores. At 15 weeks, there were minimal effects of loss of smooth muscle K_Ca_3.1 on overall lesion morphology, with no significant difference in relative stenosis (Fig. 2e), neointima area (Fig. 2f), medial area (Fig. 2h) or total vessel area (Fig. 2i). However, iSMKO mice exhibited significantly reduced lumen area (Fig. 2g). Plaque progression continued to 40 weeks at which time iSMKO mice demonstrated greater relative stenosis (Fig. 2j), neointimal area (Fig. 2k) and reduced lumen size (Fig. 2l). However, at 40 weeks, iSMKO mice showed no significant differences in medial area (Fig. 2m) or total vessel area (Fig. 2n) following smooth muscle-specific K_Ca_3.1 knockout *versus* control.

**Figure 2.**
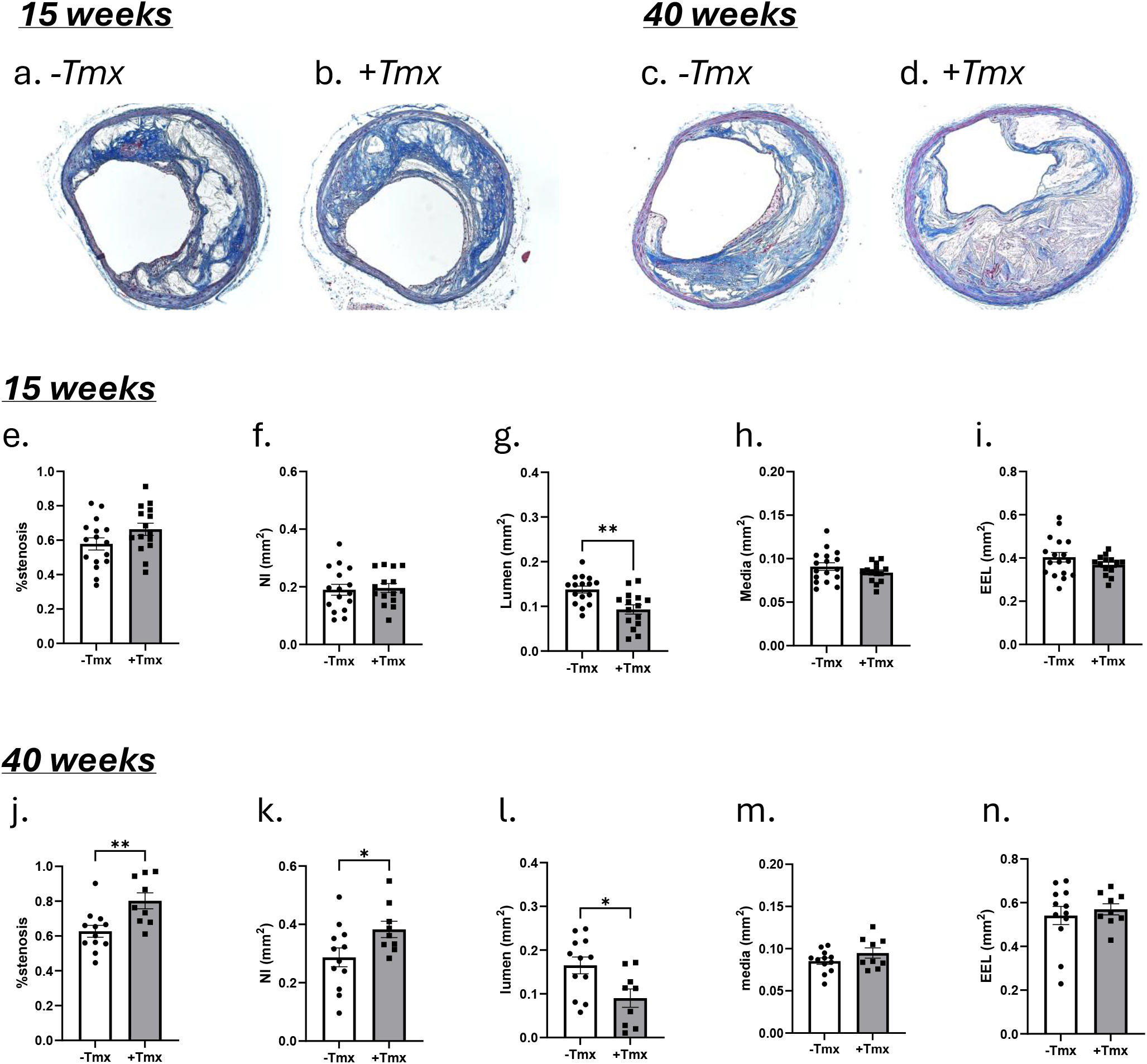
*SMC-specific K_Ca_3.1 silencing increases atherosclerotic plaque size*. Representative trichrome stained BCA sections from vehicle and tamoxifen-injected mice at 15 (a, b, resp.) and 40 weeks (c, d, resp.). Morphometric data showing relative stenosis, neointimal (NI) area, lumen area, medial area and vessel (EEL) areas at 15 (e) and 40 (f) weeks. At 15 weeks, only lumen area was significantly reduced in tamoxifen-injected mice (+Tmx, n=15) vs vehicle (-Tmx, n=16). At 40 weeks, tamoxifen-injected mice (+Tmx, n=9) showed increase relative stenosis, NI area and reduced lumen area vs. vehicle (-Tmx, n=12). *p<0.05, **p<0.01.

### Silencing smooth muscle K_Ca_3.1 alters atherosclerotic plaque composition

We and others have previously shown that inhibiting K_Ca_3.1 reduces smooth muscle cell proliferation and migration^6,7^. Furthermore, we have previously found global silencing of K_Ca_3.1 reduces smooth muscle cell content in BCA lesions^13^. Consistent with this, representative immunofluorescence images of BCA lesions from vehicle-and tamoxifen-treated mice at 15 weeks (Fig. 3a,b, respectively) and 40 weeks (Fig. 3c,d, respectively) demonstrate smooth muscle cells (SMαA+, green) and macrophages (CD68+, magenta) demonstrated reduced SMC content in lesions from iSMKO mice at both 15 and 40 weeks (Fig. 3e,j). It has been shown that the majority of SMC within atherosclerotic lesions are of medial origin and migrate into the intima during lesion development^16–18^. Previous studies by us and others ^6,7,19^ have shown the K_Ca_3.1 inhibitor, TRAM-34, inhibits SMC migration *in vitro* and reduces SMC content in atherosclerotic plaques. The effect of SMC K_Ca_3.1 silencing in reducing SMC content within the lesion is consistent with K_Ca_3.1 activation being involved in medial-to-intimal SMC migration in atherosclerosis development. Similarly, collagen content was also reduced in iSMKO mice at both time points (Fig. 3f & k). However, in contrast to a reduced macrophage content previously seen with global K_Ca_3.1 silencing^13^, lesions in iSMKO mice demonstrated greater relative macrophage content which was significant at 40 weeks. At 15 weeks, iSMKO mice showed an increase in the percentage of CD68-positive cells (Fig. 3g), which was significant when analyzed as CD68-positive area (Fig. 3h). By 40 weeks, macrophage content was further elevated, with significant increases in both the percentage of CD68-positive cells (Fig. 3l) and CD68-positive area (Fig. 3m) in tamoxifen-treated iSMKO mice compared with controls (Fig. 3e). Necrotic core area was not significantly different between groups at either 15 or 40 weeks (Fig 3i,n). Thus, overall, lesions in iSMKO animals were larger, with reduced smooth muscle and collagen content and greater macrophage infiltration. Together, these findings suggest that smooth muscle K_Ca_3.1 contributes a beneficial role in atherosclerosis by limiting lesion size and altering lesion composition by increasing smooth muscle and collagen content, while inhibiting macrophage recruitment and/or infiltration, features consistent with a smaller, more stable plaque.

**Figure 3.**
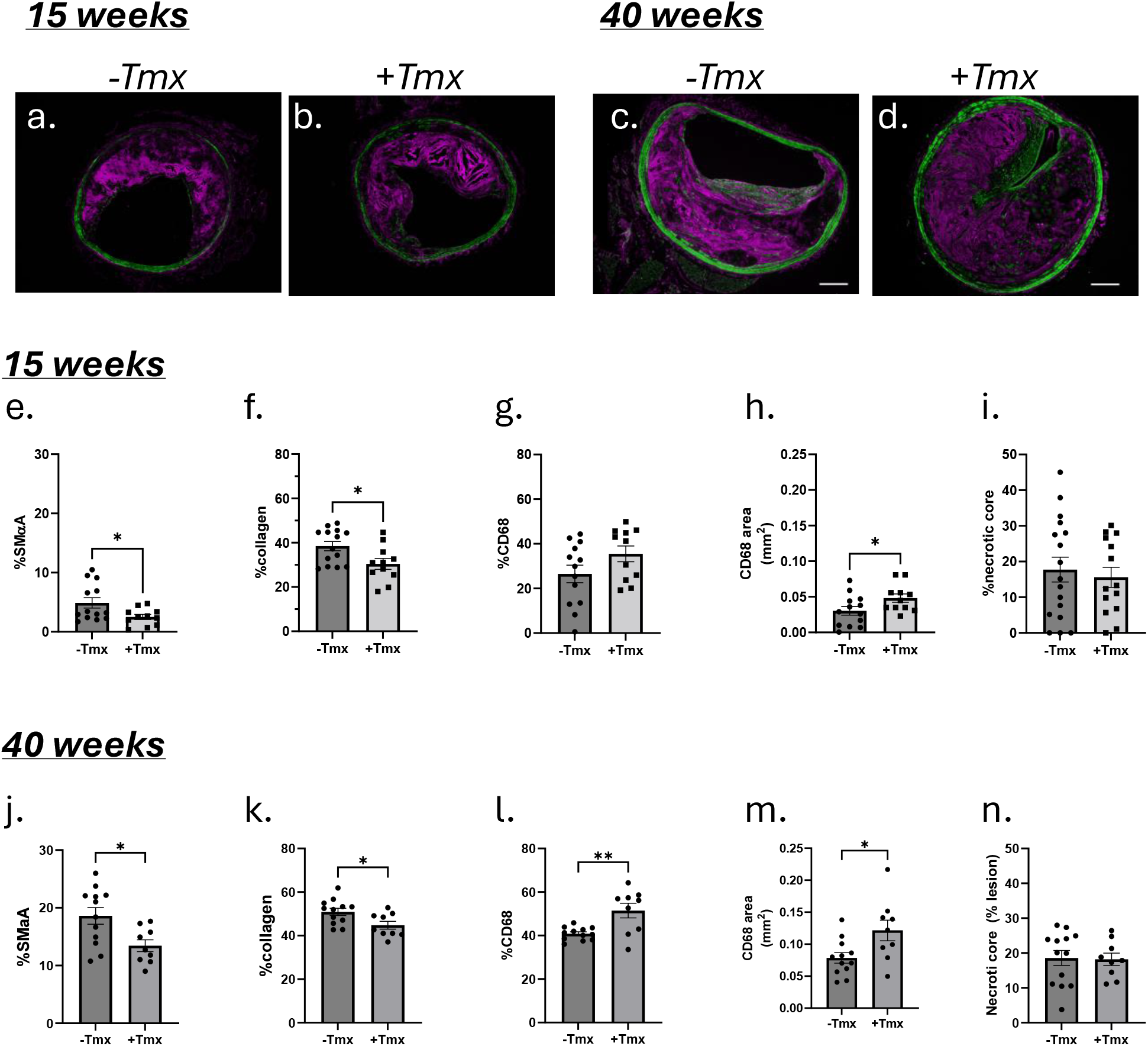
*SMC-specific K_Ca_3.1 silencing alters plaque composition*. Representative immunofluorescent BCA sections from vehicle and tamoxifen-injected mice at 15 (a, b, resp.) and 40 weeks (c, d, resp.) demonstrating smooth muscle (SMαA+, green) and macrophage (CD68+, magenta) cells. Quantitative group data of lesions showing %SMαA, %collagen, %CD68, CD68 area and %necrotic core at 15 (e) and 40 (f) weeks. At 15 weeks %SMαA and %collagen were decreased and absolute CD68 area increased in lesions of tamoxifen-injected mice (+Tmx, n=15) vs vehicle (-Tmx, n=16). These differences in lesions of tamoxifen-injected mice persisted at 40 weeks (+Tmx, n=9) with the additional increase in relative CD68 vs. vehicle (-Tmx, n=12). *p<0.05, **p<0.01.

### Transcriptome changes in atherosclerotic lesions with smooth muscle K_Ca_3.1 silencing

To explore potential novel mechanisms underlying the impact of K_Ca_3.1 silencing in smooth muscle, we performed bulk RNA sequencing of BCAs from tamoxifen-and vehicle-injected iSMKO mice prior to (baseline) and after 15 weeks on WD to identify unique transcriptomic signatures induced by K_Ca_3.1 silencing in atherosclerosis. At baseline, analysis identified 837 differentially expressed genes (DEG) significantly altered (45 downregulated and 792 upregulated, Fig. 4a). DEGs were used to identify key pathways using enrichment analysis and directional gene changes in IPA. The enriched pathway analysis identified the majority enriched pathways (8 of 10) as mitochondrial, i.e. increased oxidative phosphorylation and respiratory electron transport, and decreased mitochondrial dysfunction (Fig. 4b). Overall, these changes are very similar to those found in our prior global K_Ca_3.1 knockout study^13^. Comparison analysis within IPA confirmed this similarity between SMC-specific (iSMKO) at baseline and global (DKO) silencing with enriched mitochondrial function (Fig 4c.). Similarity predicted upstream activation of PPAR signaling, i.e. rosiglitazone and PPARGC1A, were identified by IPA (Fig 4d) at baseline in iSMKO and in DKO. Conversely, analysis of BCAs from iSMKO mice after 15 weeks of WD produced only 33 differentially expressed genes (23 down-regulated and 10 up-regulated) and pathway analysis indicated an enrichment in circadian rhythm genes and the ID3 signaling pathway and inactivation of pathways including immune cell communication (Fig 5 a,b). While evidence in the literature supports a role for some of these pathways in atherosclerosis, e.g. ID3 signaling^20^ and immune cell communication^21^, the relatively small number of significant genes that were altered lessens confidence in these as primary drivers of the changes seen in the current study. Comparison analysis in IPA of our previous data in global K_Ca_3.1 knockout (DKO), BCAs in iSMKO prior to disease development (baseline) and BCA lesions in iSMKO (Fig. 4c & d) reveal common pathway and putative upstream regulators between DKO and baseline iSMKO, that are no longer significant after WD and lesion development in iSMKO, including increased mitochondrial function, immune cell communication and rosiglitazone-associated regulation, i.e. PPAR activation.

**Figure 4.**
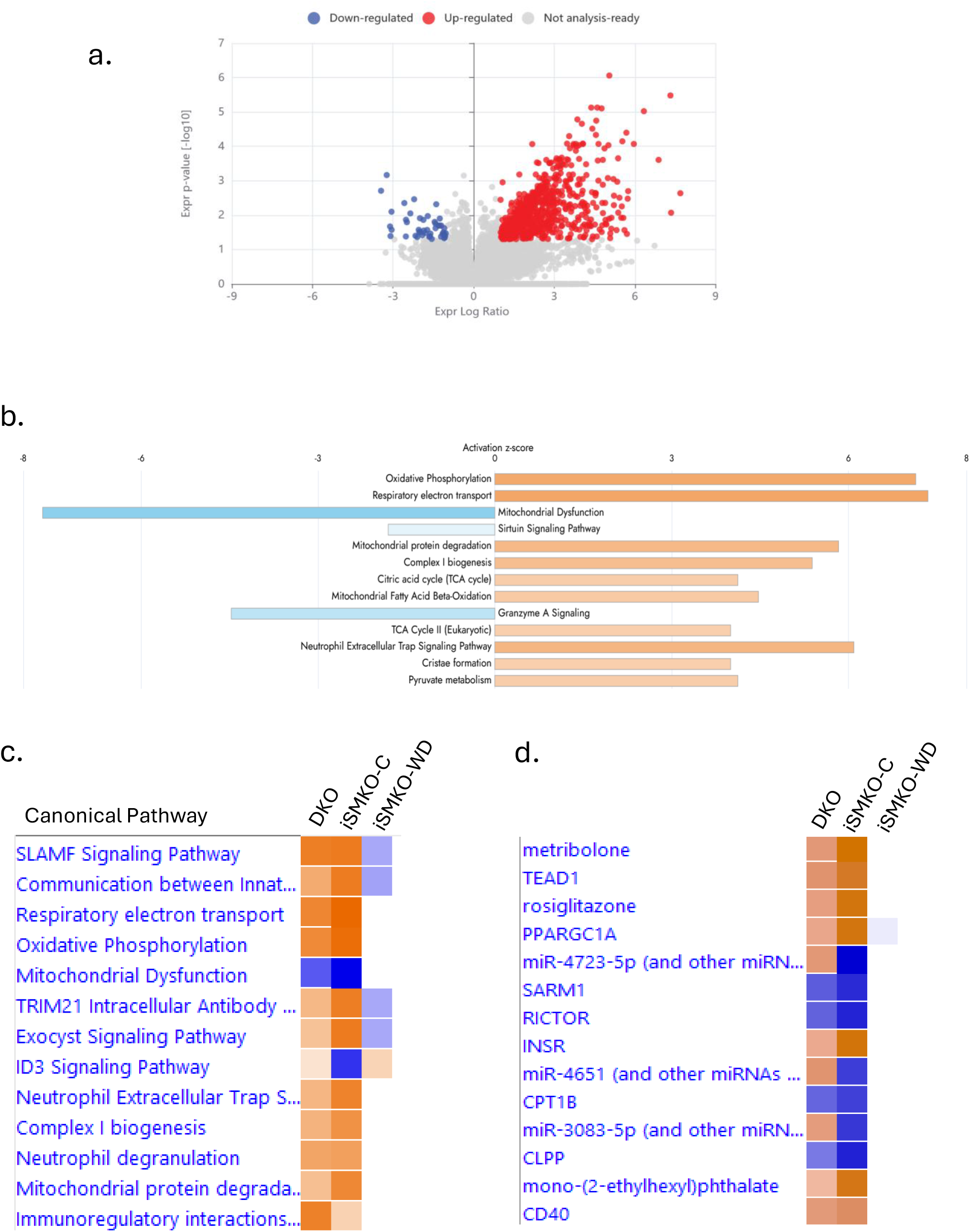
*Transcriptomic responses to smooth muscle-specific silencing of K_Ca_3.1 in BCA*. a) Volcano plot of differentially expressed genes in BCA from tamoxifen-and vehicle-injected mice on standard chow diet (baseline). Upregulated genes coded red and downregulated genes coded green (log^2^fold change <-1 and>1, adjusted p value < 0.05). b) Top 10 predicted affected pathways in IPA for +Tmx vs-Tmx demonstrating a preponderance of effect on mitochondria (8 of 10 pathways). c) Comparison analysis of significantly affected pathways in iSMKO at baseline (iSMKO-C) and after 15 weeks WD (iSMKO-WD) versus our previous findings in BCA lesions from global K_Ca_3.1 knockout mice. d) Comparison analysis of predicted upstream regulators in iSMKO at baseline (iSMKO-C) and after 15 weeks WD (iSMKO-WD) versus our previous findings in BCA lesions from global K_Ca_3.1 knockout mice.. Orange indicates predicted activation, blue indicates predicted inhibition. Overall similar changes were noted in iSMKO-C and DKO which were not present in iSMKO-WD.

**Figure 5.**
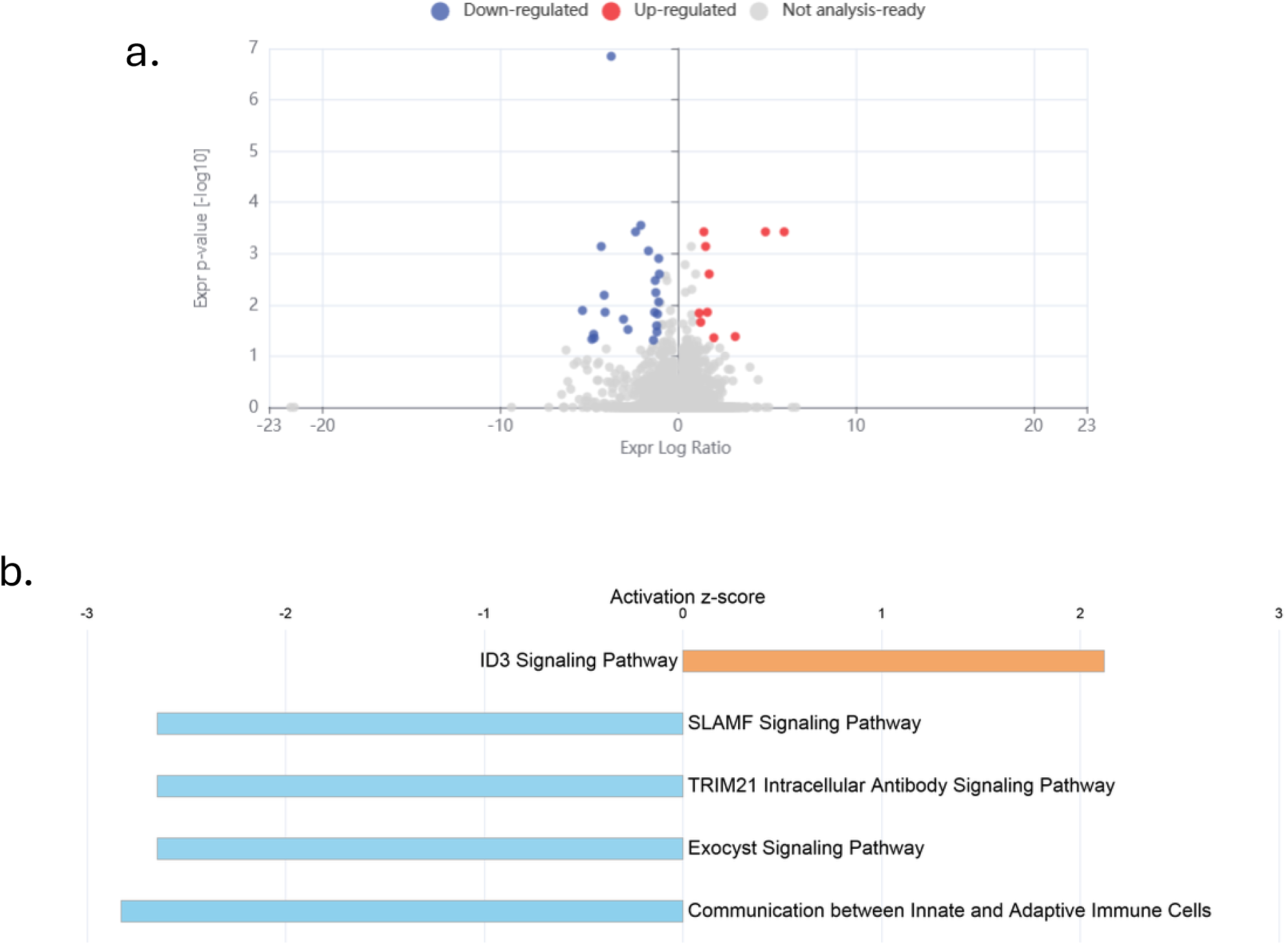

## Discussion

The current study is the first to use *in vivo* smooth muscle-specific gene silencing to examine the role of K_Ca_3.1 on atherosclerosis progression. Using Myh11-CreER^T2^ Kcnn4^fl/fl^Apoe^-/-^ (iSMKO) mice we were able to demonstrate a beneficial role of SMC K_Ca_3.1 on overall lesion size and composition in that silencing SMC K_Ca_3.1 resulted in lesions with greater neointimal size, loss of SMC and collagen content and increased macrophage content, characteristics consistent with a larger, more unstable plaque^2,22,23^. In addition, subsequent transcriptomic analysis revealed activation of similar putative beneficial mechanisms common to both global and SM-specific K_Ca_3.1 silencing at baseline, e.g. improved mitochondrial function and PPAR activation, which were subsequently lost upon disease induction.

Our finding of a reduction in SMC content in lesions is consistent with previous studies showing that both pharmacological^6,7,19^ and genetic silencing^13^ of K_Ca_3.1, inhibits smooth muscle cell proliferation and migration. Accordingly, one would expect a reduction in medial-to-intimal smooth muscle migration and expansion and secondary reduced intimal fibrosis during lesion development with K_Ca_3.1 inhibition. Consistent with this, lesions from iSMKO mice showed a reduction in BCA lesion SMC and collagen content, features associated with a more unstable plaque composition indicating a putative beneficial role of SMC K_Ca_3.1 in plaque composition.

Moreover, in contrast to a reduced macrophage content in lesions from mice with global K_Ca_3.1 knockout^13^, SMC-specific silencing K_Ca_3.1 led to an increased macrophage content as assessed by the macrophage marker, CD68. Reduced macrophage content with global silencing is consistent with previous findings that K_Ca_3.1 inhibitors reduce lesion macrophage content in Apoe^-/-^ mice^6,10^ supporting the idea of K_Ca_3.1 silencing/inhibition in macrophages as being beneficial in atherosclerosis. In macrophages, K_Ca_3.1 has been shown to regulate migration, M1/M2 polarization, respiratory burst and pathogen killing^24^. Pharmacological blockade of K_Ca_3.1 inhibits differentiation toward M1 phenotype and increases M2/M1 ratio in advanced plaques^10^. Furthermore, studies have demonstrated that K_Ca_3.1 inhibition reduces vascular inflammation^25^ and may shift macrophage polarization toward the M2 phenotype, thereby contributing to the formation of smaller, more stable plaques^26–29^.

We previously hypothesized that the reduced macrophage content with global K_Ca_3.1 silencing may be due, in part, to reduced K_Ca_3.1 in smooth muscle and loss of MCP1/Ccl2 expression and/or SMC cell transdifferentiation to foam cells. The cytokine MCP1/Ccl2 is one of several chemokines that play a key role in monocyte recruitment to atherosclerotic plaques^30^. MCP1/Ccl2 receptor-deficient mice have a substantial reduction in arterial lipid deposition^31^ and diminished numbers of macrophages in the arterial wall^32^. In addition, we previously provided evidence that K_Ca_3.1 positively regulates MCP1/Ccl2 expression in SMC but not endothelial cells^13^. However, the finding that SMC-specific silencing of K_Ca_3.1 in the current study demonstrated increased macrophage content indicates that inhibition of SMC K_Ca_3.1-mediated MCP1 does not limit macrophage recruitment/infiltration. Thus, the implication is that the reduction in macrophage content in lesions with global K_Ca_3.1 knockout is likely due to loss of K_Ca_3.1 in macrophages. Similarly, the increased macrophage content in the iSMKO mice lesions does not support a role of K_Ca_3.1-mediate SM-to-macrophage transdifferentiation which SMC lineage tracing has described to account for 50-80% of macrophage/foams cells in atherosclerotic lesions ^33^. Thus, the reduced macrophage content in lesions from mice with global K_Ca_3.1 knockout and pharmacological K_Ca_3.1 inhibition is likely due to inhibition of K_Ca_3.1 in macrophages. However, the increased macrophage content in lesions with SMC-specific K_Ca_3.1 silencing indicates some yet to be determined protective effect of smooth muscle K_Ca_3.1 on macrophage recruitment/retention in plaques.

Loss of K_Ca_3.1 in smooth muscle resulted in overall detrimental effects on plaque size and composition indicative of a beneficial role of smooth muscle K_Ca_3.1 in atherosclerosis. Overall, transcriptomic analysis of iSMKO prior to WD feeding (baseline) identified a similar transcriptomic signature in BCAs to that observed previously with global knockout^13^. Specifically, improved mitochondrial function and PPAR pathway activation emerged as top putative mechanisms underlying the effects of both global and SMC-specific K_Ca_3.1 silencing. The decreased SMC content of lesions in iSMKO mice is consistent with the known role of K_Ca_3.1 inhibition on SMC proliferation and migration. Accordingly, both putative downstream mediators of K_Ca_3.1 inhibition, i.e. improved mitochondrial function and PPAR activation, are also known to inhibit SMC proliferation and migration. Increased mitochondrial respiration and reduced mitochondrial dysfunction have been consistently associated with decreased atherosclerotic progression^34–39^. In smooth muscle, mitochondrial dysfunction has been associated with increased migration and phenotype modulation^34,39,40^. Similarly, studies have shown that PPAR agonists reduce smooth muscle cell migration and proliferation^41^ in addition to reducing vascular inflammation, plaque size, and atherosclerosis progression^42–45^,. Thus, the similarity of transcriptomic changes in global and SMC-specific K_Ca_3.1 loss suggests a key role for mitochondrial function and PPAR signaling in mediating effects on SMC during atherosclerosis, likely explaining the reduction in SMC content in lesions of both global and SMC-specific knockout mice. However, a key difference between global and SMC-specific K_Ca_3.1 silencing in atherosclerosis is that while both lesion size and macrophage content were lower with global KO^13^, both were conversely increased in iSMKO lesions. One conclusion is that these aspects of lesion development were affected by loss of K_Ca_3.1 in non-SMCs, such as macrophages and/or endothelial cells. K_Ca_3.1 is known to be expressed in macrophages and activation increases M1 to M2 transition and cytokine production ^9,10^ and pharmacological inhibition and gene silencing of K_Ca_3.1 has been shown to reduce macrophage infiltration/content, lipid content and intimal area of atherosclerotic lesions resulting in lesions with characteristics of more stable plaques^6,10,13^. However, the cell type(s) responsible for these beneficial effects of loss of K_Ca_3.1 in global KO remain to be determined. The relative contribution of K_Ca_3.1 among cell types will require cell-specific, inducible silencing/overexpression and, potentially, fate tracking to elucidate responsible cell type(s). Comparison of transcriptomic changes in lesions from iSMKO and global K_Ca_3.1 KO failed to provide tangible differences, as significant transcriptomic changes were lost upon WD-feeding and lesion formation. Given that K_Ca_3.1 silencing was restricted to smooth muscle, the relative increase in non-SM cells retaining K_Ca_3.1, e.g. endothelial cells, fibroblasts, macrophages, T-cells, B-cells, and platelets, may account for the loss of differential expression from baseline to 15-week WD and lesion formation. Determination of the cell type(s) responsible for the remaining effects of K_Ca_3.1 inhibition will require cell-specific silencing of other cell types and/or single cell transcriptomics.

In conclusion, our study provides further support for the important contribution of K_Ca_3.1 activation in the progression of atherosclerotic lesion development and composition and provides novel insights into cell-specific contributions of K_Ca_3.1. While previous studies using pharmacological or global genetic silencing of K_Ca_3.1 have shown an overall beneficial effect on atherosclerotic lesion size and macrophage infiltration, SMC-specific loss of K_Ca_3.1 demonstrates an increase of lesion size and macrophage content, while retaining the reduction in SMC content. Thus, the beneficial vs. detrimental effects of K_Ca_3.1 inhibition in atherosclerosis are cell-specific, involving a potentially beneficial role of SMC K_Ca_3.1 in modulating lesion size and composition with a commitment detrimental role of K_Ca_3.1 activation in other cells types, e.g. macrophages. Thus, while pharmacological, global inhibition of K_Ca_3.1 may be a viable avenue for limiting overall lesion size, there are underlying potentially detrimental effects on SMC content which may negatively alter plaque stability and overall outcome. Further refinement of the effects of K_Ca_3.1 in other cell types on lesion development *in vivo* will be necessary for refinement of cell specific targeting of K_Ca_3.1 in therapies for clinical benefit.

## Data Availability

The data that support the findings of this study are available from the corresponding author, DKB, upon reasonable request.

## Author contributions

P.A. contributed to the experimental design, collected and analyzed data and wrote the first draft of the manuscript; D.L.T contributed to the conception and experimental design and contributed to final editing of the manuscript; H.J.B contributed to RNAseq analysis and final editing of the manuscript; H.B. collected and analyzed data and contributed to final editing of the manuscript; S.B.B contributed to the analysis interpretation and final editing of the manuscript and D.K.B. contributed to the conception and experimental design and final editing of the manuscript

