## Supplemental Methods for "Smooth muscle K_Ca_3.1 limits plaque progression and alters plaque composition in mice"

**Supplement- Methods**

*Generation of SM-specific $K_{Ca}3.1$, ApoE knockout mice*. We generated conditional, SM-specific $K_{Ca}3.1$ knockout mice using Kcnn4$^{fl/fl}$ mice generated from ES cells with loxP sites flanking exon 4 of Kcnn4 (Biocytogen, Waltham, MA; Figure x). CRISPR/Cas9 gRNAs was used to delete exon 4 of Apoe in Kcnn4$^{fl/fl}$ mice in the MU Animal Modeling Core (University of Missouri, Columbia, MO). Kcnn4$^{fl/fl}$,Apoe$^{-/-}$ mice which were subsequently crossed with Myh11-CreER$^{T2}$ mice (Jax:019079) to produce Myh11-CreER$^{T2}$ Kcnn4$^{fl/fl}$Apoe$^{-/-}$ mice (iSMKO). Mice were housed in identical cages in the same room with 12-hour light-dark cycles and controlled temperature and humidity withing the Veterinary Medicine building. Only male mice were used for this study due to the Y-chromosome insertion of the Myh11-CreER$^{T2}$ transgene. At 8-10 weeks of age, mice were administered tamoxifen (1 mg i.p.) or vehicle daily for 5 days to selectively delete $K_{Ca}3.1$ in smooth muscle. Mice then switched to a Western diet (0.2% cholesterol, 42% kcal from fat; TD.88137, Teklad Diets) for 15 or 40 weeks. Mice were anesthetized, and either perfusion-fixed via the left ventricle at 80 mmHg with 10% neutral-buffered formalin for histological analysis, or the brachiocephalic artery (BCA) was carefully dissected and rapidly frozen in liquid nitrogen for molecular analysis. The BCA from each animal was then dissected and stored in formalin until it was embedded in paraffin for subsequent histology.

*Immunofluorescent.* Immunofluorescence was performed as previously described[4]. Sections were incubated with avidin–biotin two-step blocking solution (Vector SP-2001) to inhibit background staining and 3% hydrogen peroxide to inhibit endogenous peroxidase. Non-serum protein block (Dako X909) was then applied to inhibit non-specific protein binding. Sections were incubated at 4°C overnight with primary antibodies SMαA (Dako, #M0851), or CD68 (Invitrogen, #MA513324). After washing, sections were incubated with biotinylated secondary antibody in phosphate-buffered saline containing 15 mM sodium azide and peroxidase-labeled streptavidin (Dako LSAB+ kit, peroxidase, K0690). Diaminobenzidine (DAB, Dako) was applied 5 minutes for visualization of the reaction product, sections were then counterstained with haematoxylin, dehydrated, and coverslipped. Images of the sections were obtained using an Olympus BX61 photomicroscope and Spot Insight Color camera (Diagnostic Instruments). The relative area and mean density of positive staining were determined for each section of interest utilizing ImagePro Plus (Media Cybernetics).

*Morphology and immunohistochemistry*: Serial sections were obtained from the BCA and stained using Verhoeff-Van Gieson (VVG) and Masons Trichrome as previously described[4].

Morphometric measurements were obtained on spatially calibrated images of VVG stained sections using standard planimetry and NIH Image J software (Bethesda, MD). Intimal area was defined as the area between the internal elastic lamina (IEL) and the luminal border of the artery while the area between the IEL and external-elastic lamina (EEL) was referred to as the medial area. Intima-media ratio (I/M) was calculated as the ratio of intimal- over medial-areas. Percent stenosis was defined as intimal area/IEL area. Relative intimal collagen content was determined from trichrome images using the blue channel following color deconvolution using Fiji (Image J). Necrotic core (NC) was defined as acellular areas within a lesion that were negative for VVG staining (i.e., extracellular matrix was lacking with total or almost complete loss of collagen). Boundary lines were delineated around those regions and the area was measured by image analysis software as described above. Based on the criterion by Seimon et. al. [46], areas <3,000 $\mu m^2$ were not counted as they likely do not constitute substantial areas of necrosis. Relative NC area was calculated by dividing total NC area by total lesion area. All analyses were conducted by a single investigator blinded to the experimental group.

*Blood analysis*: Total cholesterol content was determined by IDEXX BioAnalytics (Columbia, MO) from serum collected the day of euthanasia.

*RNA isolation:* RNA isolation from BCA was performed by using the RNeasy Plus Mini Kit (Qiagen Catalog # 74136) following the standard protocol. Briefly, sample lysis buffer was added to the BCA and aorta tissue samples. Then, BigPrep Lysing Matrix D was added to each sample, and the tubes were placed in a FastPrep homogenizer. The samples were homogenized for 20-30 seconds at maximum speed (6.0 m/s) to ensure complete tissue disruption. The process was repeated as needed to achieve a homogeneous sample for further analysis. After homogenization, the samples were passed through a gDNA column to remove genomic DNA. The flow-through was then mixed with an equal volume of 70% ethanol. The mixture was subsequently applied to an RNA elution column, followed by washing steps according to the kit's instructions. RNA was then eluted, and the concentration and quality of the RNA were assessed using a Nanodrop spectrophotometer. The same RNA isolation protocol was followed for the *in vitro* cell culture experiments, except for the FastPrep homogenization step.

*Transcriptomic analysis*. For transcriptomic analysis, BCAs were snap-frozen to create samples from both groups (n= 6-9 per group). Sequencing was conducted by the MU Genomics

Core. To generate Illumina Stranded mRNA libraries, the poly-A containing mRNA was purified from total RNA, RNA was fragmented, double-stranded cDNA was generated from fragmented RNA, and the unique dual index containing adapters were ligated to the fragment ends. All libraries generated were sequenced together on an Illumina NovaSeq flow cell using paired-end read, 100 base read length sequencing. Total yield was ~2 billion reads providing ~50 million read pairs per sample. Analysis and visualization of the NGS sequence data was performed by the MU Informatics Research Core Facility (IRCF).

RNA-Seq data were processed and analyzed as previously described[47]. Briefly, latent Illumina adapter sequence was identified and removed from input 100-mer RNA-Seq data using Cutadapt. Subsequently, input RNA-Seq reads were trimmed and filtered to remove low quality nucleotide calls and whole reads, respectively, using the Fastx-Toolkit. To generate the final set of quality-controlled RNA-Seq reads, foreign or undesirable sequences were removed by similarity matching to the Phi-X genome (NC_001422.1), the relevant ribosomal RNA genes as downloaded from the National Center for Biotechnology Information, or repeat elements in RepBase, using Bowtie. The remaining RNA-seq reads were aligned to the Mus musculus genome assembly GRCm39 using the Bioconductor package Rsubread with default settings for paired reads. Rsubread uses a “seed-and-vote” mapping paradigm which allows for quick mapping of subreads for gene expression analysis. This provided the initial gene expression estimates which were annotated according to their Entrez ID number using the Bioconductor database org.Mm.eg.db. These initial expression estimates were then normalized and sorted into a comparison matrix based on genotype using the Bioconductor package DESeq2. The two comparison groups, Apoe-/- and DKO, were then analyzed according to their differential gene expressions with DKO used as the treatment group to be compared against the Apoe-/- expressions. Outlier samples were identified through the use of Principal Component Analysis (PCA) and Counts Per Million plots. Normalization and differential gene expression (DGE) analysis was then rerun after the exclusion of aforementioned outliers. The final results were written into an excel file and visualized using PCA and volcano plots, highlighting genes with an absolute log2 fold change (|log2FC|) >2 and adjusted p-values <0.05. Genes with the lowest adjusted p-values were analyzed using DAVID (david.abcc.ncifcrf.gov)[48] and IPA (Ingenuity Systems) as previous[49-54].
